# A systematic review and standardized clinical validity assessment of male infertility genes

**DOI:** 10.1101/425553

**Authors:** Manon. S. Oud, Ludmila Volozonoka, Roos M. Smits, Lisenka E.L.M. Vissers, Liliana Ramos, Joris A. Veltman

**Affiliations:** Department of Human Genetics, Donders Institute for Brain, Cognition and Behavior, Radboud University Medical Centre, 6500 HB, Nijmegen, The Netherlands; Institute of Genetic Medicine, Newcastle University, NE1 3BZ, Newcastle, United Kingdom; Scientific Laboratory of Molecular Genetics, Riga Stradins University, LV-1007, Riga, Latvia; Department of Obstetrics and Gynecology, Division of Reproductive Medicine, Radboud University Medical Centre, 6500 HB, Nijmegen, The Netherlands

**Keywords:** Male infertility, spermatogenic failure, genetics, clinical validity, gene-disease relation, gene panel, next-generation sequencing, systematic review

## Abstract

**Study question:** Which genes are confidently linked to human male infertility?

**Summary answer:** Our systematic literature search and clinical validity assessment reveals that a total of 67 genes are currently confidently linked to 81 human male infertility phenotypes.

**What is known already:** The discovery of novel male infertility genes is rapidly accelerating with the availability of Next-Generation Sequencing methods, but the quality of evidence for gene-disease relationships varies greatly. In order to improve genetic research, diagnostics and counseling, there is a need for an evidence-based overview of the currently known genes.

**Study design, size, duration:** We performed a systematic literature search and evidence assessment for all publications in Pubmed until June 2018 covering genetic causes of male infertility and/or defective male genitourinary development.

**Participants/materials, setting, methods:** Two independent reviewers conducted the literature search and included papers on the monogenic causes of human male infertility and excluded papers on genetic association or risk factors, karyotype anomalies and/or copy number variations affecting multiple genes. Next, the quality and the extent of all evidence supporting selected genes was weighed by a standardized scoring method and used to determine the clinical validity of each gene-disease relationship as expressed by the following six categories: no evidence, limited, moderate, strong, definitive or unable to classify.

**Main results and the role of chance:** From a total of 23,031 records, we included 1,286 publications about monogenic causes of male infertility leading to a list of 471 gene-disease relationships. The clinical validity of these gene-disease relationships varied widely and ranged from definitive (n=36) to strong (n=12), moderate (n=33), limited (n=86) or no evidence (n=154). A total of 150 gene-disease relationships could not be classified.

**Limitations, reasons for caution:** Our literature search was limited to Pubmed.

**Wider implications of the findings:** The comprehensive overview will aid researchers and clinicians in the field to establish gene lists for diagnostic screening using validated gene-disease criteria and identify gaps in our knowledge of male infertility. For future studies, the authors discuss the relevant and important international guidelines regarding research related to gene discovery and provide specific recommendations to the field of male infertility.

**Study funding/competing interest(s):** This work was supported by a VICI grant from The Netherlands Organisation for Scientific Research (918-15-667 to JAV).

## Introduction

### Introduction

Infertility, a common disorder with a world-wide prevalence affecting 15% of all couples in the reproductive age, is defined as the inability to conceive within one year of unprotected sexual intercourse (Zegers-Hochschild et al. 2009). It is suggested that approximately 7% of the male population is affected by a factor of infertility and that these collectively explain half of all infertile couples (Krausz and Riera-Escamilla 2018; Irvine 1998; Winters and Walsh 2014).

The etiology of infertility is highly heterogeneous, which is not surprising when considering that both male and female reproductive systems need to function in a combined and precisely coordinated fashion in order to conceive a child. Appropriate genetic regulation is one of the most important and indispensable prerequisites to control the coordination and timing of sexual development and fertility. Because of the complexity of the gamete development, the interference of a genetic origin is suspected. Studies aiming to elucidate the genetic basis of fertility defects in both human and mice have defined numerous crucial pathways for male infertility, including sexual differentiation, development of the genitourinary system and gametogenesis (Krausz and Riera-Escamilla 2018; Jamsai and O’Bryan 2011). Currently more than 900 male infertility genes have been described in the Jackson Laboratory’s Mouse Genome Informatics (MGI) database (http://www.informatics.jax.org/), and 2,300 testis-enriched genes are currently known in human (Schultz, Hamra, and Garbers 2003).

### Genetic testing in infertility

It is currently thought that at least 15% of all human male infertility patients can be explained by genetic defects (Krausz and Riera-Escamilla 2018). Since the discovery of an extra X chromosome in Klinefelter patients (47,XXY) as the first genetic cause of infertility in the late 1950’s (Ferguson-Smith et al. 1957; Jacobs and Strong 1959), more than 3,500 papers have been published on the genetics of male infertility, implicating various common genetic origins as well as hundreds of other genes in male infertility. Despite these large numbers, genetic diagnostic testing is usually confined to karyotyping, AZoospermia Factor (AZF) deletion screening and Cystic Fibrosis Transmembrane conductance Regulator (*CFTR*) mutation analysis and leaves a vast majority of patients unexplained. Currently a genetic diagnosis is reached in about 4% of all infertile males - a number that has not increased since the late 1990’s (Tuttelmann, Ruckert, and Ropke 2018; Johnson 1998). This is in sharp contrast to the increase in diagnostic yield seen for other conditions with a strong genetic component and increase in large-scale technologies for genetic testing since then (Rehm 2017; Tuttelmann, Ruckert, and Ropke 2018). Importantly, the lack of a genetic diagnosis limits clinicians in providing personalized information about the potential success of Assisted Reproductive Technologies (ART), resulting in many couples undergoing these invasive procedures such as TEsticular Sperm Extraction (TESE), without any chance of success. ART may lead to a situation where infertility becomes an inherited condition. Therefore, a lack of genetic diagnosis limits counseling for couples involved with regard to the reproductive health of offspring that can be conceived by ART (Belva et al. 2016).

The diagnostic yield for genetic testing in male infertility remains low for several reasons. Firstly, the condition is highly heterogeneous and thousands of genes are thought to play a role in spermatogenesis (Schultz, Hamra, and Garbers 2003). High-impact mutations in any of these genes will always remain at very low frequency in the population because of their impact on fitness. This means that in order to find recurrently mutated genes and confidently linked novel genes to infertility, one has to screen large cohorts of patients for pathogenic variants in large numbers of genes. This has been laborious and expensive for a long time due to limitations of traditional genetic assays such as Sanger sequencing. Since the first introduction of Next-Generation Sequencing (NGS) in 2005, the technology has evolved to allow rapid and affordable sequencing of large amounts of DNA (Metzker 2010). This has expedited sequencing of large gene panels, all coding genes (the exome) and even whole genomes (Payne et al. 2018). In contrast to other fields in medical genetics and oncology where NGS has revolutionized disease gene identification as well as genomic diagnostics, the use of NGS in male infertility has only recently commenced and its use in routine diagnostics is still very limited.

The second reason underlying a disappointingly low diagnostic yield for male infertility is that the interpretation of genetic data is hampered by gaps in our understanding of the biology of male spermatogenesis and (in)fertility. This urgent need for better understanding of cellular, molecular biochemical and genetic mechanism(s) is highlighted by a recent study of the World Health Organization, who listed this need as one of the key areas of research focus (Barratt et al. 2017).

### Clinical validity assessment of gene-disease relationships

With the introduction and advances in genomics, the number of genes associated to male infertility has expanded in recent years. However, the amount of genes confidently linked to disease is still very limited in comparison to developments in other genetic diseases such as intellectual disability (Tuttelmann, Ruckert, and Ropke 2018; Vissers, Gilissen, and Veltman 2016). This is caused in part by a lack of solid evidence linking variation in individual genes to human male infertility. The notion of sub-optimal quality of evidence in male infertility research is not limited to genetic studies but is considered a general concern in the field of reproductive biology (Barratt 2016; Evers 2013; Glujovsky et al. 2016).

In order to robustly link gene dysfunction to disease, one needs to consider multiple levels of evidence. This is especially important since insufficient, inconclusive and low-quality evidence may result in incorrect and misleading conclusions about gene-disease relationships. Moreover, if this wrongful gene-disease relation is not identified and corrected, it may lead to inappropriate diagnoses and even mismanagement and counseling of infertile couples involved. Furthermore, these incorrectly characterized genes may complicate follow-up research by contaminating candidate disease gene lists and pathway analyses.

To evaluate genetic variant pathogenicity, standard guidelines such as those provided by the American College of Medical Genetics and Genomics (ACMG) are invaluable (Richards et al. 2015). Their well-established framework combines variant allele frequency in control populations, computational prediction programs such as SIFT (Kumar, Henikoff, and Ng 2009) and PolyPhen (Adzhubei et al. 2010), as well as functional evidence and variant segregation evidence to classify sequencing variants into (likely) benign, pathogenic or variants of uncertain significance (VUS). These guidelines, however, have been developed for diagnostic purposes and are based on the assumption that the causal link between the gene and the condition has already been established. So whereas individual genetic variants may be considered pathogenic, unfortunately, the majority of candidate genes in male infertility still have questionable evidence and cannot be confidently linked to human disease.

Recently, the Clinical Genome Resource (ClinGen) has developed an extensive framework to assess the clinical validity of a gene-disease relationship (Strande et al. 2017). However, the overall number of validated disease genes is currently very limited (n=333) and does not contain any genes involved in male infertility. Another, more simplified and pragmatic version of this framework was recently published to more easily assess the clinical validity of gene-disease relationships (Smith et al. 2017). In this study, we applied this latter gene-disease scoring system to curate all available information on the genetics of human male infertility from 1958 up to June 2018. This analysis allowed us to objectively classify the evidence for the involvement of genes in male infertility as non-existing, limited, moderate, strong or definitive. The results from this work may be useful in both research and diagnostics, for example for developing diagnostic gene panels and hopefully help to strengthen genetic research in male infertility.

## Materials and methods

### Search strategy and study selection

Two independent reviewers conducted a literature search in Pubmed according to the PRISMA guidelines (Moher et al. 2009) for English articles in peer-reviewed journals. The search was performed on several occasions with the last search taking place on the 21^st^ of June 2018 without further restrictions on publication date. The search query and screening strategy aimed to collect all records of genetics research in defective male reproductive development and function (Supplemental Table S1). We excluded all papers that did not describe human patients. Since the scope of our review is limited to monogenic causes of male infertility and/or defective male genitourinary development, we excluded papers describing chromosomal aneuploidies, complex chromosomal rearrangement or copy number variations not attributable to a single gene, polygenic and multifactorial causes, as well as variants that are associated with infertility, but do not directly influence gene function such as SNP or genome-wide association studies. We also excluded genetic disorders causing severe syndromic forms of infertility, affecting multiple organ systems (in addition to the reproductive system). This for instance excludes syndromes which compromise viability, or cause physical or intellectual disabilities to such a degree that patients are unlikely to seek for help to reproduce (Supplemental Table S1).

We included patients with delayed puberty, completely sex-reversed individuals with male phenotype (46,XX maleness) and male patients with partial virilization and a Prader score of 4, or more. We used reviews on the genetics of human male infertility to supplement our strategy with papers that were not identified in our systematic search, but did report potential gene-disease relationships. Publication inclusions of doubt were resolved by discussion and consensus between all authors.

### Data extraction and assessment of clinical validity

From eligible papers presenting original data, we extracted the gene names, patient phenotypes, inheritance pattern, method of discovery and whether or not single nucleotide or copy number variants were identified in the genes mentioned in infertile men. After extraction of the gene names from all records, we employed a recently published gene-disease scoring system to establish the strength of evidence for the relationship between a gene and male infertility (Smith et al. 2017).

A detailed description of the evidence assessment and a assessment template are described in Supplemental Table S2 and S3. In short, for each gene, we collected evidence for the most likely mode of inheritance (recessive, dominant, X-linked, Y-linked) of the infertility (sub)phenotype primarily based on evidence provided in the original papers and from model organisms. If the human mode of inheritance was unclear, we used computational methods based on statistical learning to predict the most likely mode of inheritance (Quinodoz et al. 2017; Lek et al. 2016). All variants described were re-classified using the standard ACMG guidelines for the interpretation of sequence variants (Richards et al. 2015). Only patients who had a variant(s) that 1) match the expected or proved inheritance pattern of the disease and 2) were classified as “Pathogenic”, “Likely pathogenic” or “Uncertain significance” were eligible for scoring.

Except for variants in *CFTR* which may be more common due to founder effects in North European populations (Bombieri, Seia, and Castellani 2015), we used a maximum allele frequency of 1% in the general population as a threshold value. Variants causing fully penetrant monogenic severe male infertility suffer from strong selection in the general population and are unlikely to reach higher allele frequencies than 1% (Eilbeck, Quinlan, and Yandell 2017). Variants that were more common were classified as (likely) benign. Next to various freely available population databases such as GnomAD (Lek et al. 2016), we also used an anonymized local database with exome variants found in 3,347 fathers of children who have been referred for trio Whole Exome Sequencing (WES) in the Radboud University Medical Centre. The healthy fathers reflect the general Dutch population and to our knowledge conceived naturally.

In order to award points for statistical evidence in autosomal recessive (AR) forms of infertility described in families, we used the Logarithm of the ODds (LOD) scores from the original paper. If no LOD score was given, we used a simplified formula as provided by the Clinical Genome Resource Gene Curation Working Group (Strande et al. 2017). For dominant/X-linked diseases we used: 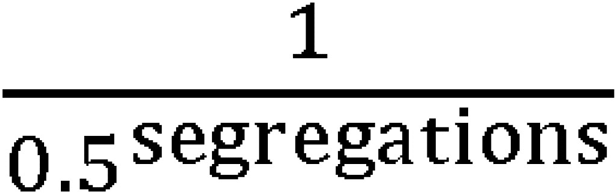 and for recessive diseases we used: 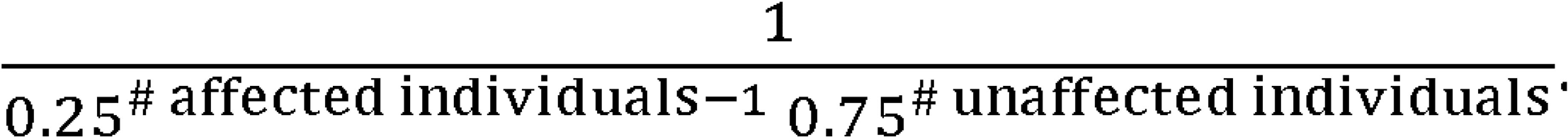 For gene function, disruption, protein interaction and model organism evidence, we critically reviewed available information from the original articles, cited papers, other (more recent) papers from Pubmed, the Protein Atlas database(Uhlen et al. 2015), the STRING database (Szklarczyk et al. 2017) and the Mouse Genome Informatics database (Smith et al. 2018). The first paper describing a variant in a potential disease gene was used as the index patient. In line with this, points for independent publications were only given from the second publication on describing variants in the same gene in unrelated patients. We then calculated the sum of the assigned points for each gene and determined the clinical validity category according to the original method (Smith et al. 2017). All genes received a denomination based on the points gathered; 1-2 points: “No evidence”; 3-8 points: “Limited”; 9-12: “Moderate”; 13-15: “Strong”; 16-17 points: “Definitive”. Similar to the publication selection process, disagreements and debatable cases were solved by consensus between all authors.

In order to prevent bias in gene-disease evaluation, a second and a third reviewer independently reviewed and verified a random selection 12 and 16 gene-disease relationships, respectively. A maximum difference of 1 point per gene-disease relationship was allowed if the classification was not altered. All other cases were discussed and re-evaluated after consensus was obtained.

### Overview of biological knowledge

From all genes with at least limited evidence, we also extracted I) the reported or expected results of semen analysis (if available), II) whether the patients described are sporadic or familial cases, and III) whether the type of infertility was isolated, a reproductive organ syndrome, endocrine disorder or part of another syndrome. All genes with at least limited evidence were plotted according to their biological function.

## Results

### Search strategy and study selection

With our search strategy, we aimed to identify all publications covering the genetics of male infertility, including those underlying syndromes affecting the endocrine system, disorders of sex development and genitourinary anomalies. Our search yielded a grand total of 23,031 publications that date from 1958 - 2018. Based on title and abstract, 18,095 studies were excluded because the publication was not in written English, or the study topic did not match our inclusion criteria (Supplemental Table S1). Although severe syndromes including male infertility phenotypes were excluded because affected patients are unlikely to seek for help to reproduce because of severe physical or intellectual disabilities, we included milder syndromes and syndromes affecting the reproductive organs only. A total of 4,936 publications were left. Since the scope of our systematic review is monogenic male infertility, we then excluded papers based on full-text screening which described genetic association or risk factors (n=668), AZF deletions (n=469), CNVs affecting multiple genes (n=28) or chromosomal anomalies (n=1,180). In addition, we excluded 803 publications that, based on full-text analysis in retrospect, were not covering the topic of the genetics of male infertility and we excluded 41 papers of which the full text was unavailable. We then screened the reference lists from included reviews (n=576) and were able to add another 115 publications that were not identified by our search strategy. In total, our search yielded 1,286 publications that met our inclusion criteria (Figure 1).

**Figure 1:**
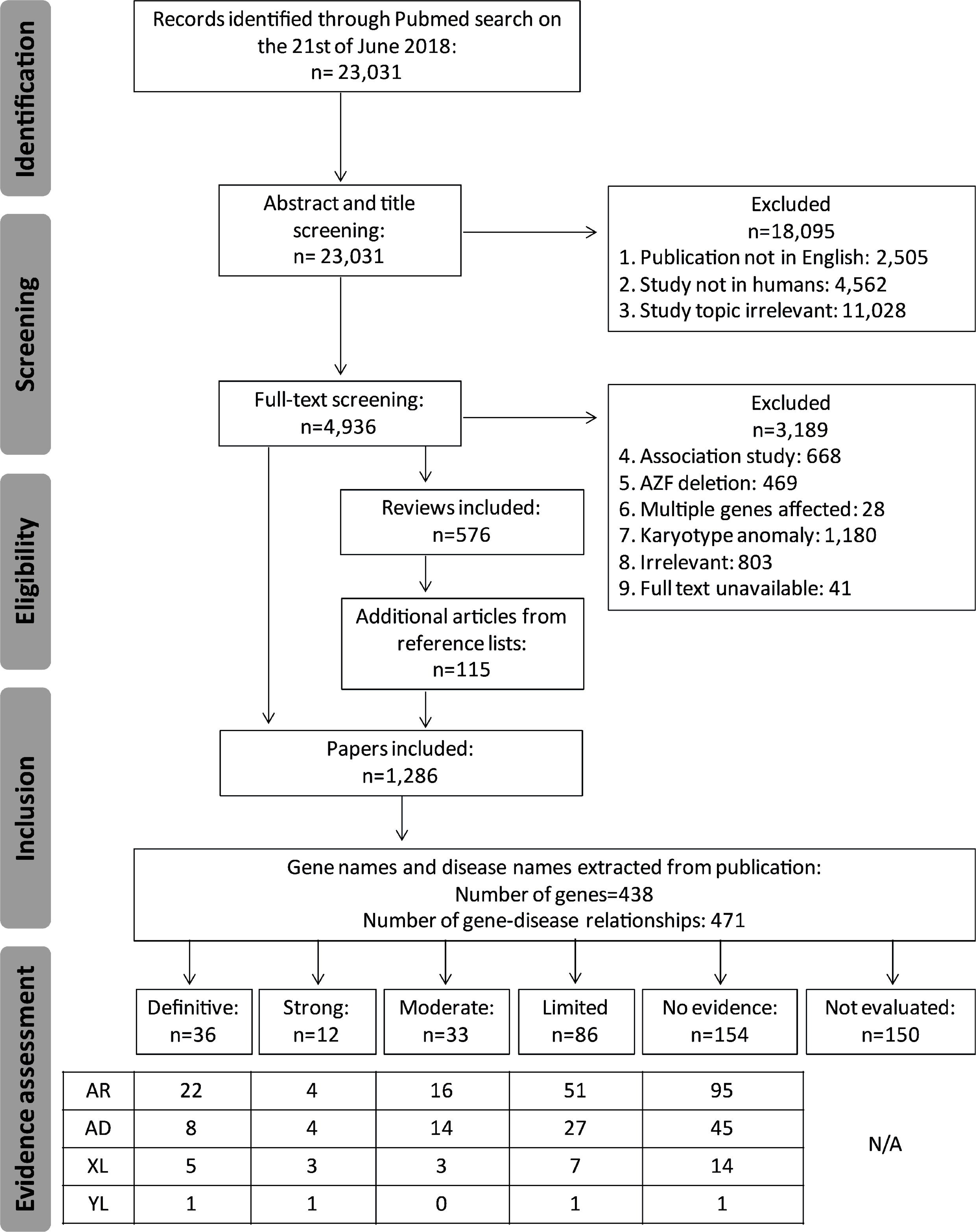
PRISMA flow chart. Our search and screening strategy to identify publications and genes eligible for clinical validity assessment. AR: Autosomal Recessive; AD: Autosomal Dominant; XL: X-linked; YL: Y-linked

The systematic literature search revealed a total of 150-200 publications per year in the past 10 years and showed that the majority of publications from the last few years report on monogenic causes of male infertility (46% in 2017), followed by genetic association or risk factor analysis (28% in 2017) (Figure 2A and B). Furthermore, the absolute number of karyotype studies has been relatively stable over the past 20 years at approximately 30 publications per year.

**Figure 2:**
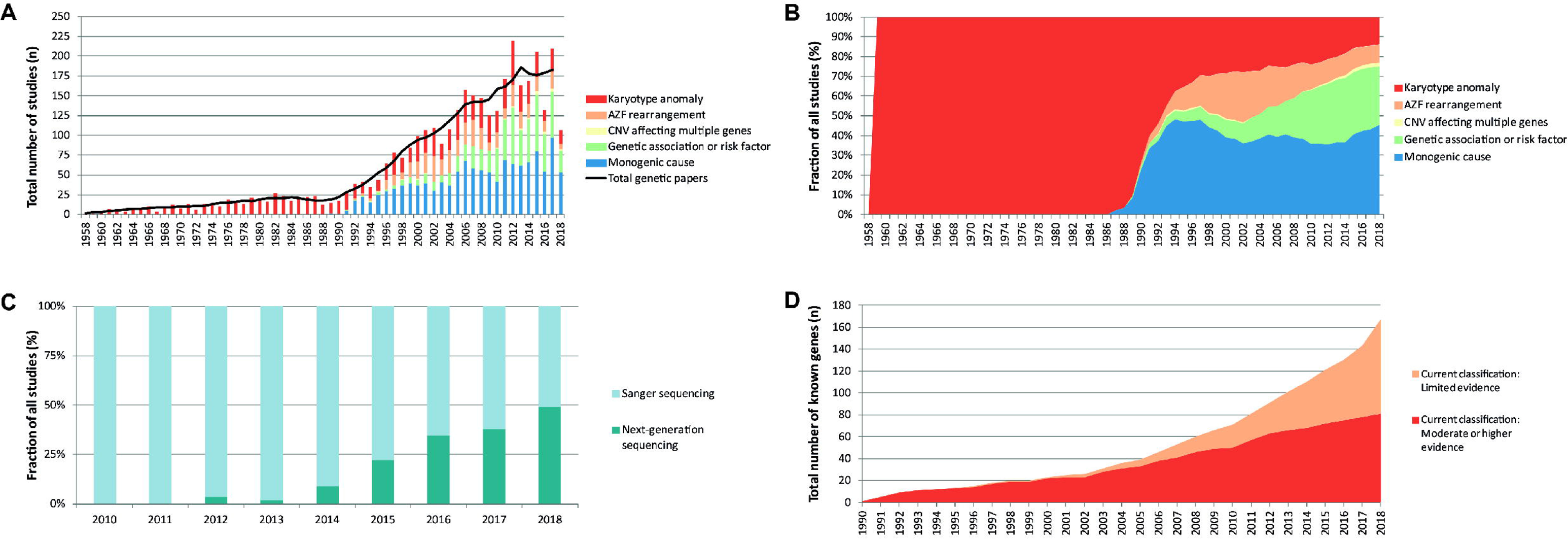
Genetic studies in male infertility. A) Graphical overview of genetic studies in male infertility. B) Graphical representation of type of genetic research in male infertility. C) The use of Sanger Sequencing and Next Generation Sequencing for the discovery of genes in male infertility. D) Increase of genes linked to human male infertility.

### Data extraction and evaluation of evidence

From the 1,286 included publications, we extracted 438 unique HUGO approved gene names and 471 gene-disease relationships (Figure 1). The number of gene-disease relationships is higher than the number of genes because several genes were described in multiple male infertility phenotypes. A further look into the discovery method of these gene-disease relationships showed that DNA sequencing has been the most commonly used technique for novel gene discovery and replication studies (84% of all publications). At the moment a shift from Sanger sequencing to Next-Generation Sequencing methods is taking place (Figure 2C).

We then assessed the clinical validity of each gene-disease relationship by using the simplified scoring system designed to establish the strength of a relationship between a single gene and a Mendelian disease (Smith et al. 2017) (Supplemental Table S2 and S3). In short, the scoring system takes into account the total of unrelated patients, the number of papers that reproduced the initial finding, the number of unique pathogenic variants and the evidence of gene disruption by the variant and the phenotype of model organisms.

After excluding genes that did not contain any potentially pathogenic variant or were unable to be classified, a total of 173 gene-disease relationships were curated and classified into the following categories definitive (n=36), strong (n=12), moderate (n=33), limited (n=86) and no evidence (n=6). We identified a total of 67 genes that can at least be moderately linked to a total of 81 male infertility or abnormal genitourinary development phenotypes showing autosomal recessive (n=42), autosomal dominant (n=26), X-linked (n=11) and Y-linked (n=2) inheritance patterns. Patients were found sporadic (n=15), in families (n=10) or in both (n=56) and led to either isolated (n=18), reproductive organ or endrocrine syndrome (n=49) or a syndromic form of infertility (n=14). A summary of the results is depicted in Table 1 and Supplemental Table S4; full scoring is available in Supplemental Table S5. In 154 cases, no (likely) pathogenic variants were identified and were therefore also classified as “No evidence” for involvement in human infertility without further curation of other evidence (Supplemental Table S6). In 150 cases, we could not evaluate the gene-disease relationship because either the inheritance pattern remains unclear or suggests polygenic inheritance, the technical quality of the identification method was too poor or the exact variant information could not be retrieved (Supplemental Table S7).

**Table 1:**
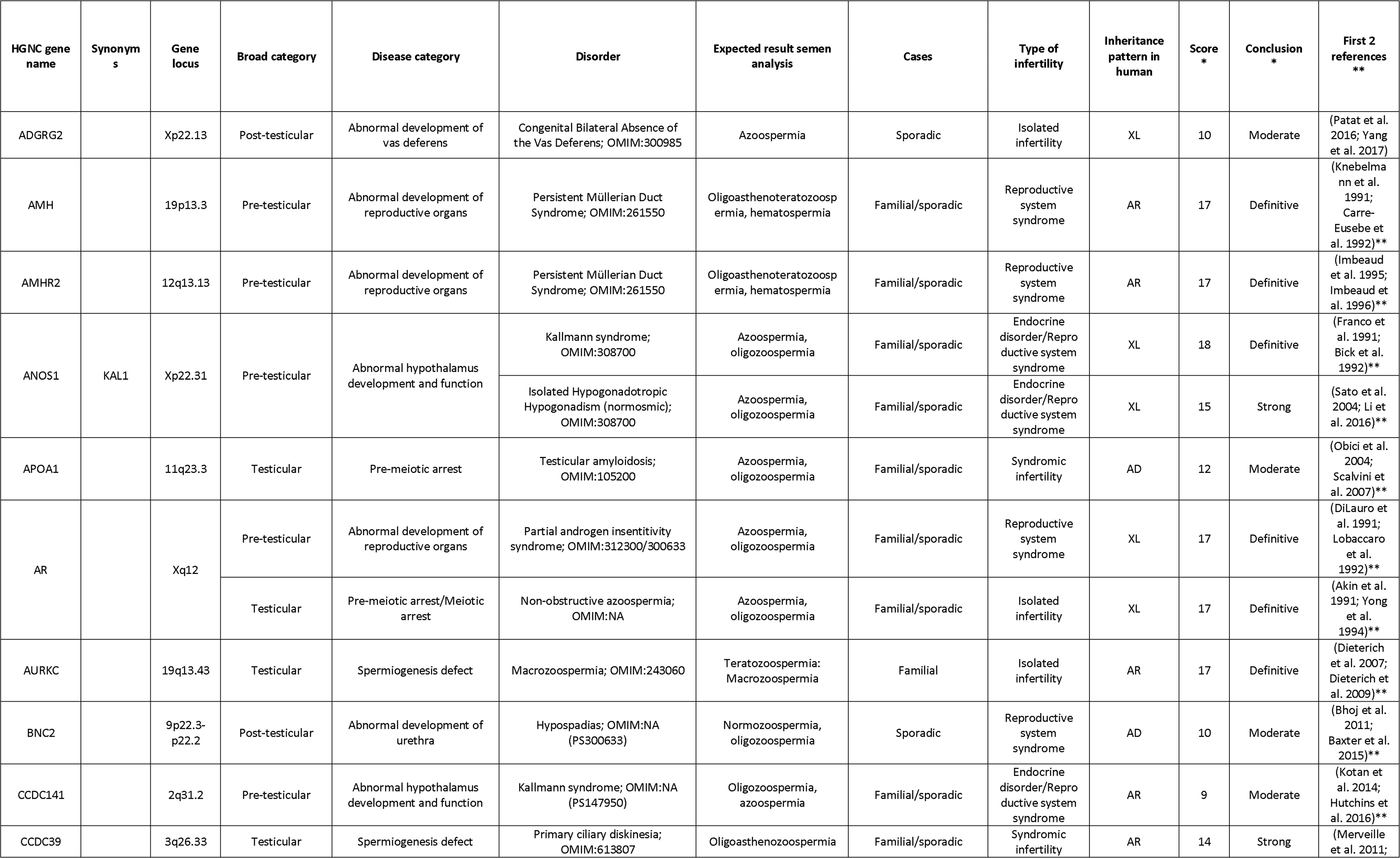
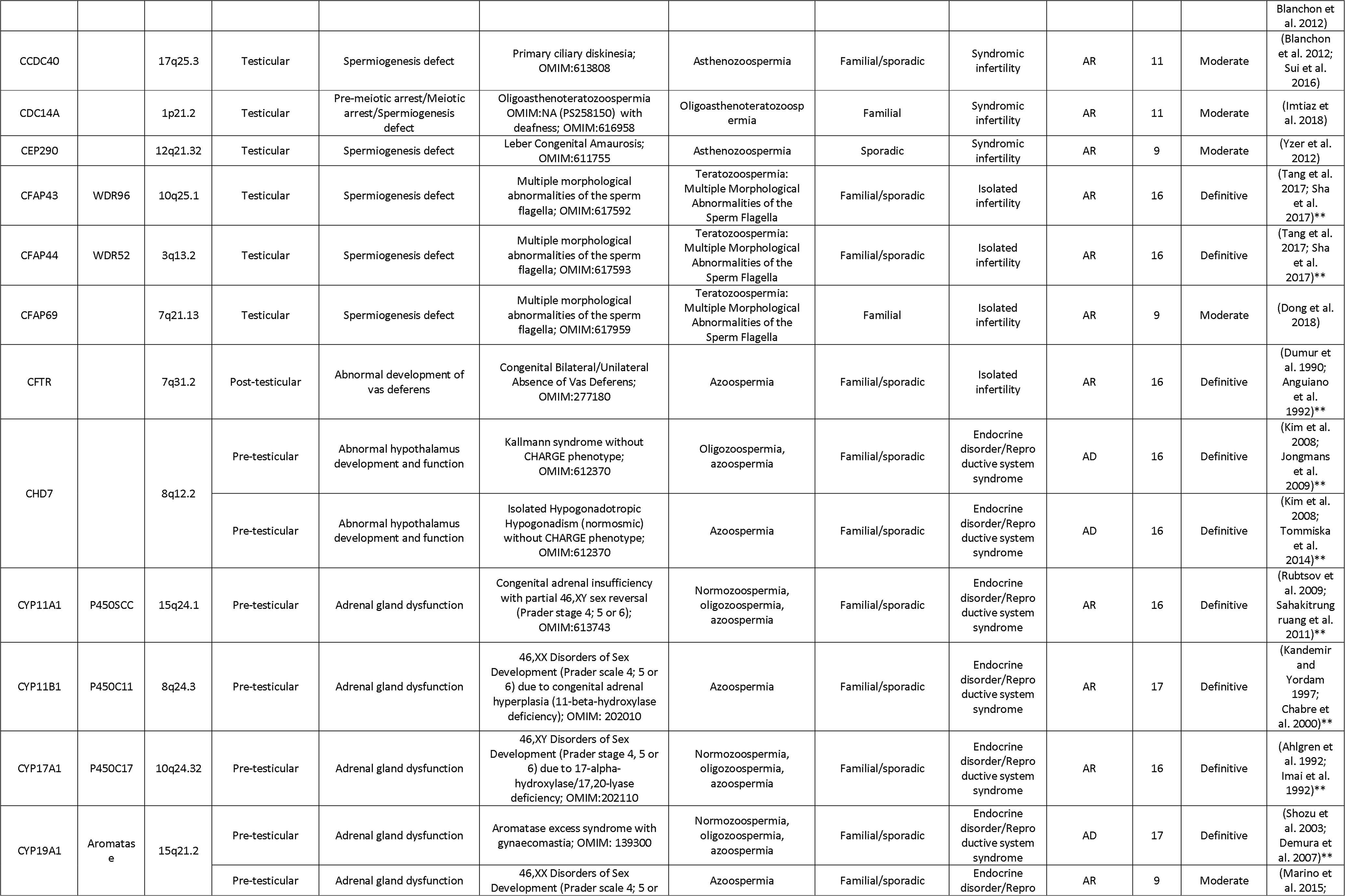
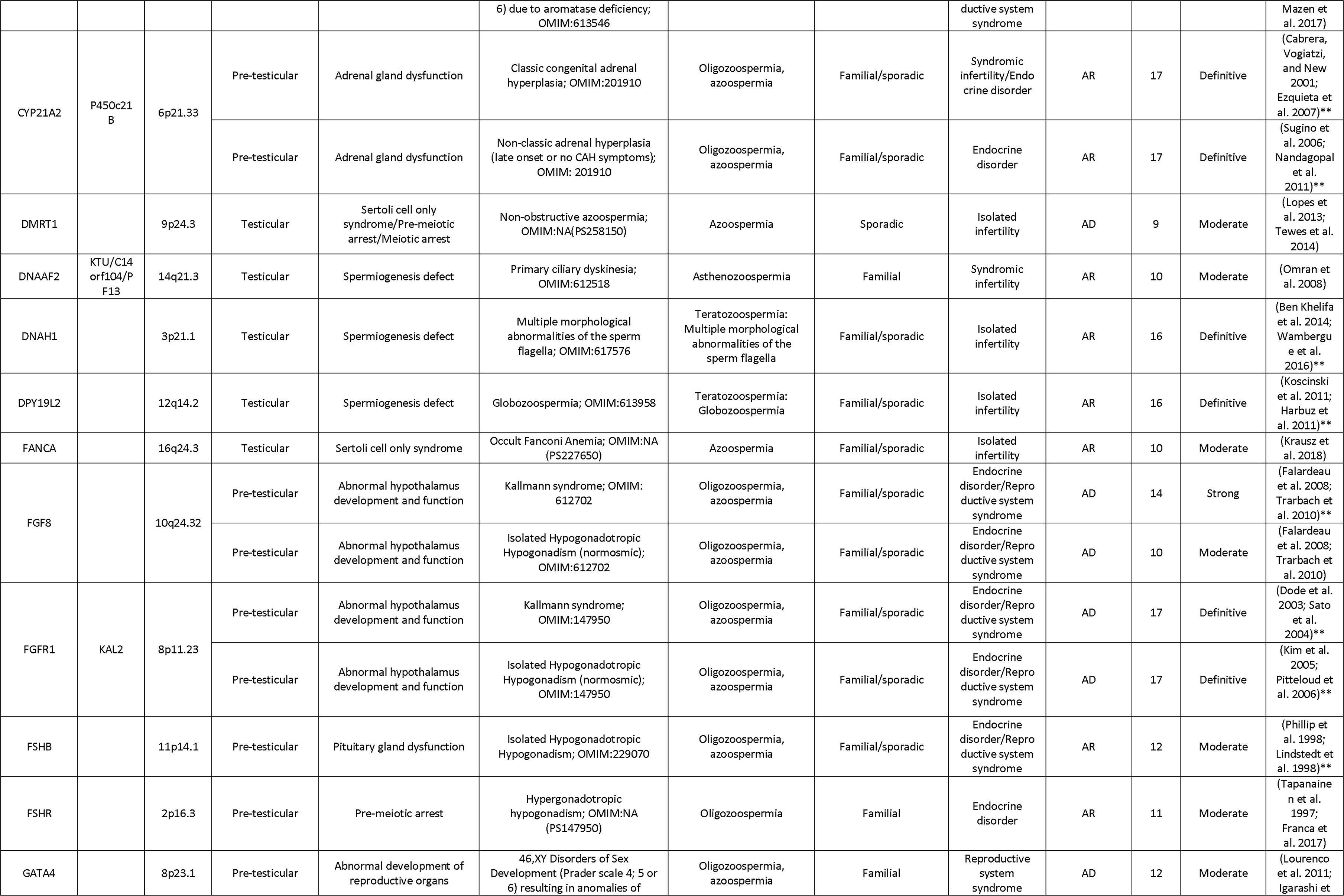
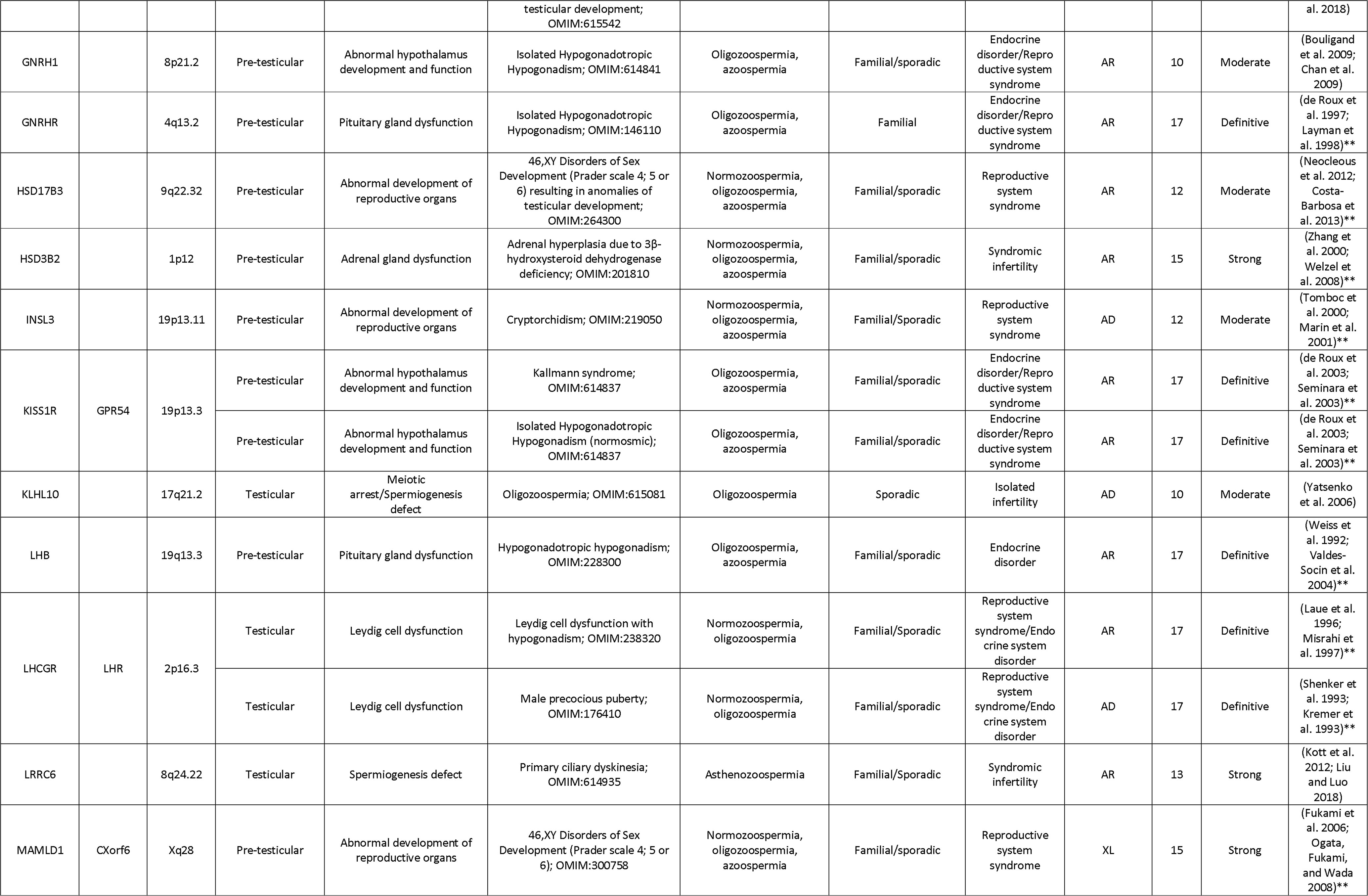
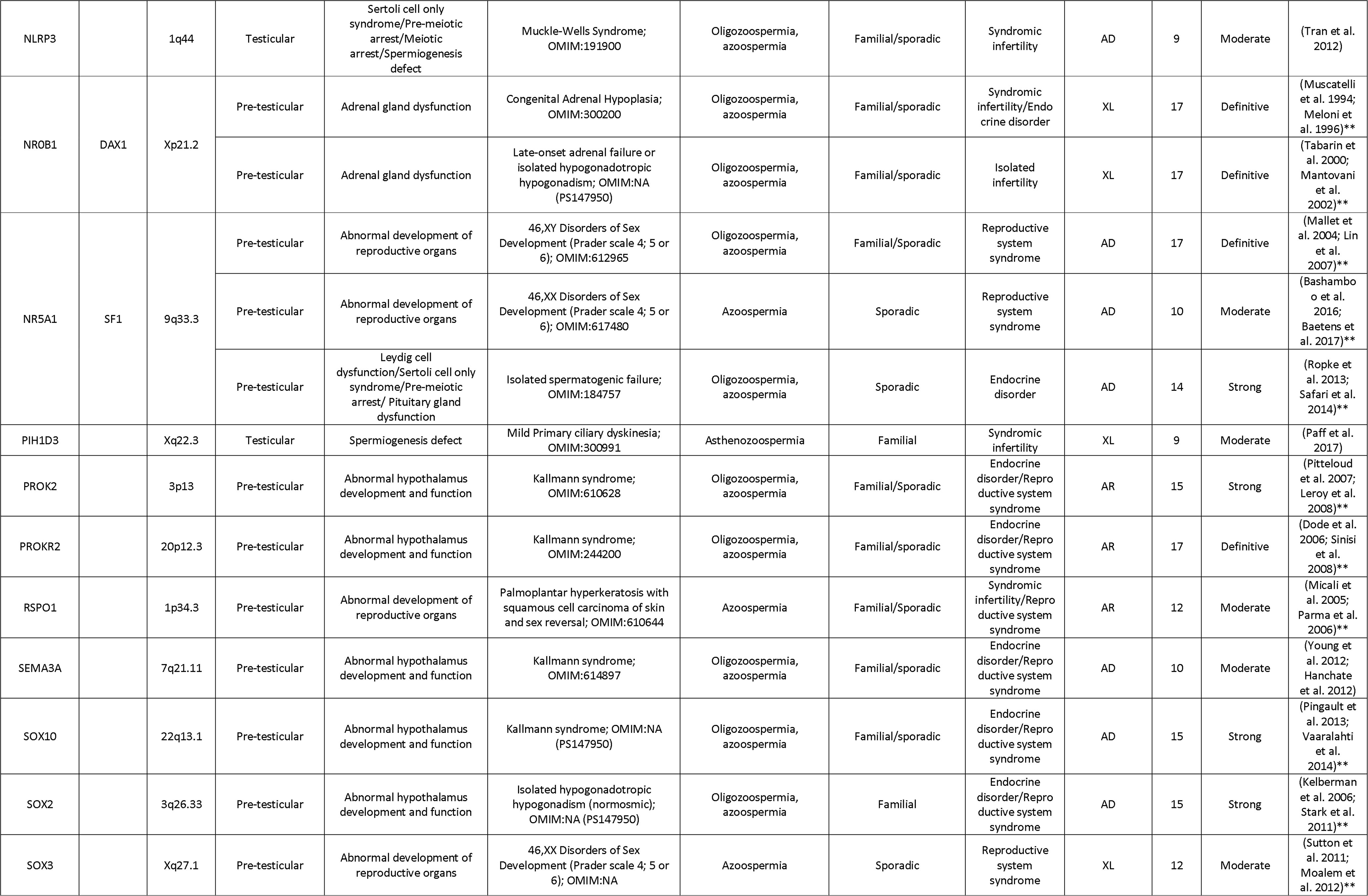
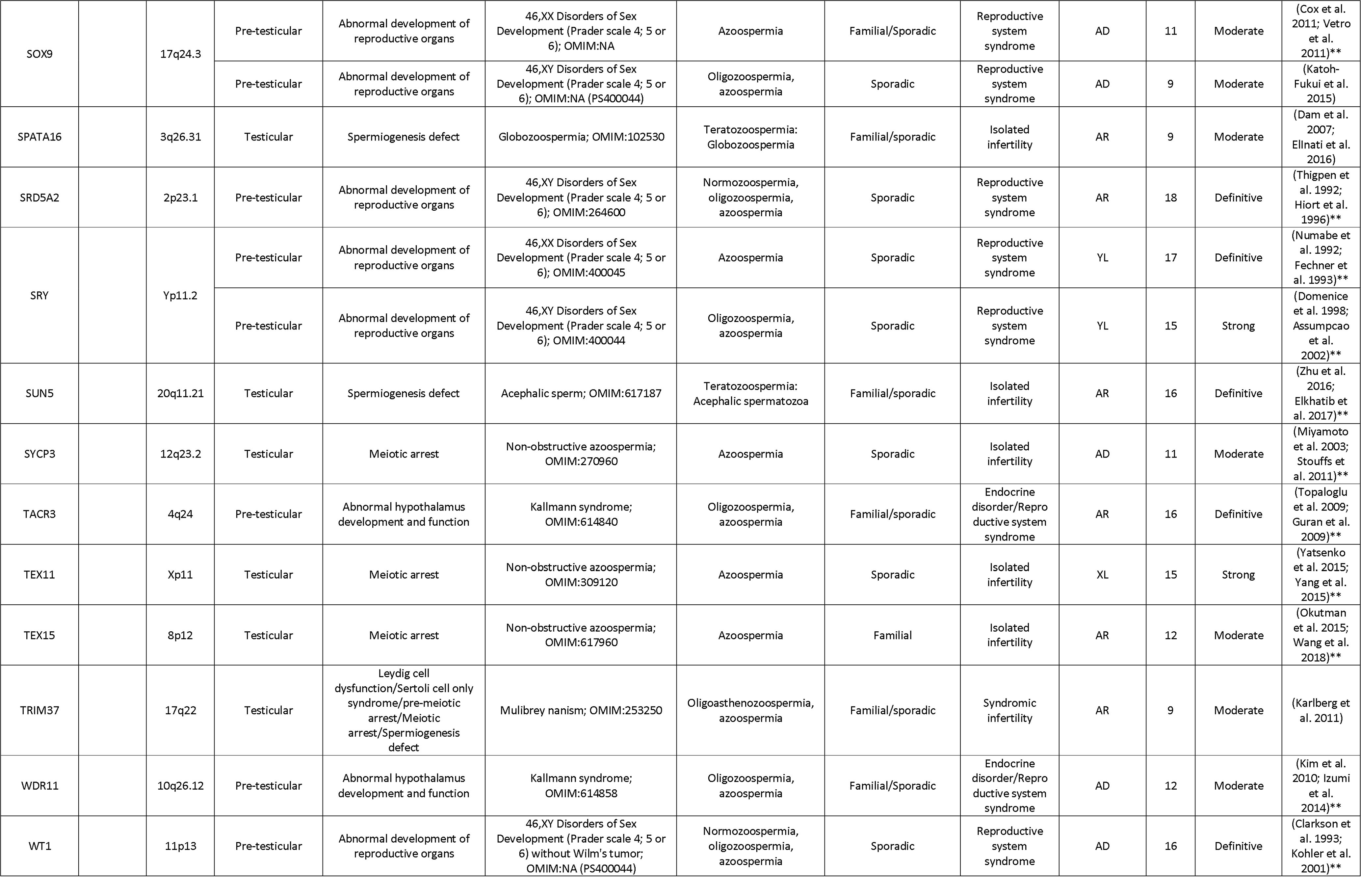
Results gene-disease relationships; only showing at least moderate evidence.

Abbreviations: HGNC: HUGO Gene Nomenclature Committee; OMIM: Online Mendelian Inheritance in Man; PS: Phenotype Series; AR: Aut osomal Recessive; AD: Autosomal Dominant; XL: X-linked; YL: Y-linked

Full table including gene-disease relationships with “Limited evidence” and “No evidence” available in Supplemental Table S4.

*Details about the score available in Supplemental Table S5.

**Additional references are available in Supplemental Table S5

The results show that the total number of confidently linked genes is growing steadily at about 3 genes per year (Figure 2D). The increase of NGS methods being used has caused an exponential growth in novel candidate genes. However, the vast majority of these are currently classified as “Limited evidence”.

### Overview of human genes involved in human male infertility

Taking into account that normal functioning of the male reproductive system is biologically mostly dictated by the hypothalamic–pituitary–gonadal axis functioning, the origins of male infertility can be divided in three major groups: pre-testicular, testicular and post-testicular. We grouped all genes with at least limited evidence for an involvement in human male infertility into these three groups based on their reported biological function (Figure 3) to assess whether the curated genes play a role in these biological processes.

**Figure 3:**
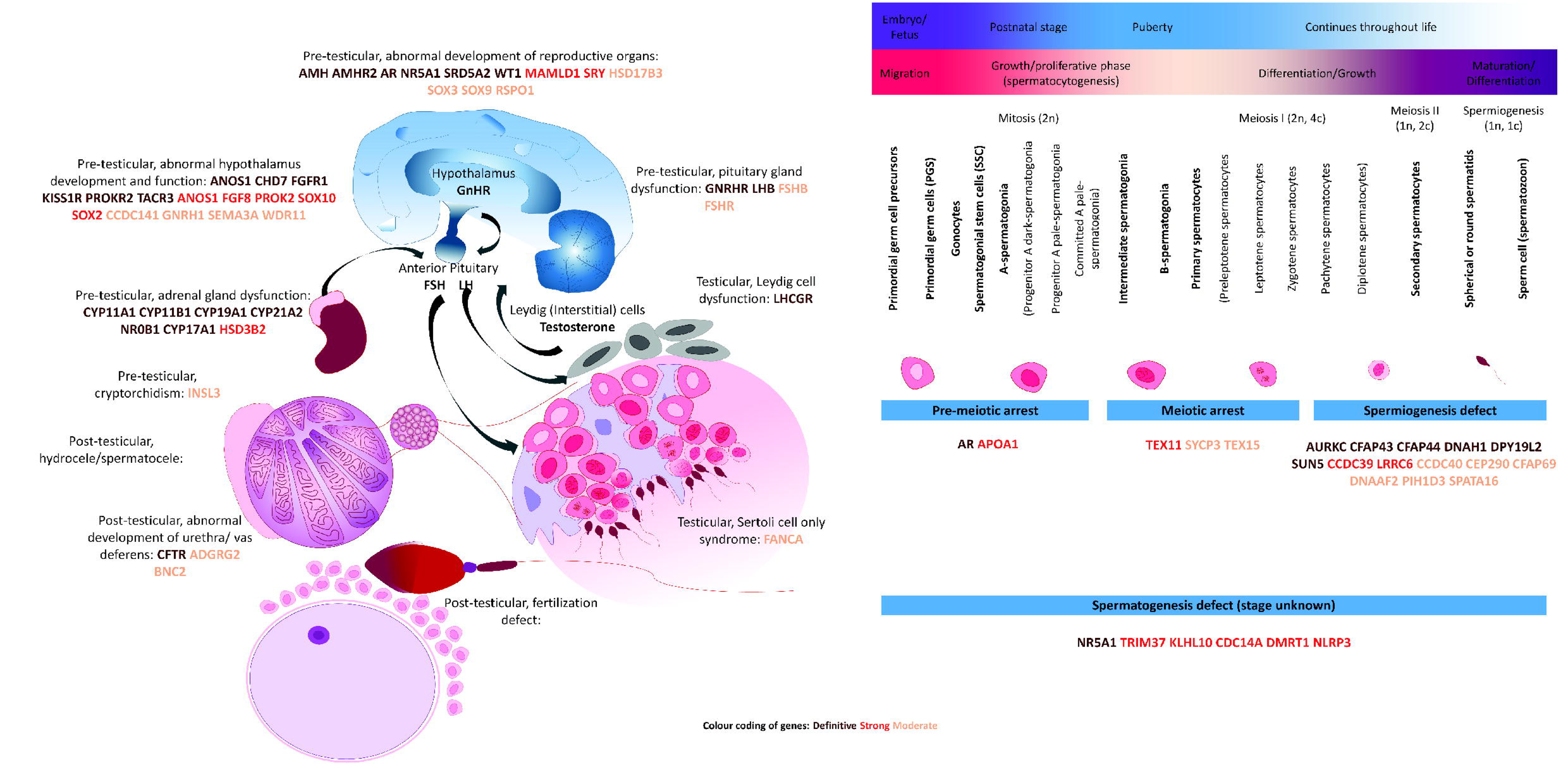
Biological overview of the genetics of male infertility. The color of each gene indicates the amount of evidence: Brown: Definitive; Red: Strong; Orange: Moderate.

Our results show that pre-testicular forms of infertility are mostly syndromic and caused by endocrine abnormalities, characterized by low levels of sex steroids and abnormal gonadotropin levels. Post-testicular causes include ejaculatory disorders or obstructions, which impair the transport of spermatozoa from the testis. These obstructions can be caused by a congenital unilateral or bilateral absence of the vas deferens. The most common genetic cause of obstructive azoospermia (OA) are biallelic variants in the *CFTR* gene (Anguiano et al. 1992; Culard et al. 1994; Dumur et al. 1990; Oates and Amos 1994; Patrizio et al. 1993) and variants in the recently identified X-linked gene *ADGRG2* (Patat et al. 2016).

Despite the fact that monomorphic forms of teratozoospermia are extremely rare, the majority of genes known to cause isolated testicular forms of infertility are involved in such disorder (n=8, 53% of all 15). The number of genes confidently linked to oligozoospermia or azoospermia when mutated remains limited (n=7, 47% of all 15).

## Discussion

Male infertility is a complex multifactorial condition which pathogenesis can be explained by environmental causes, urological conditions such as retrograde ejaculation, defective endocrine control of spermatogenesis such as hypogonadotropic hypogonadism or by the occurrence of genetic alteration in genes important for proper reproductive functioning. This standardized clinical validity assessment focused on the genetic causes of infertility and provides a systematic and comprehensive overview of all genes implicated as a monogenic cause of male infertility. Our study aimed to provide an overview of all currently available evidence and gene-disease relationships, as well as formulate a set of recommendations for future studies involving the genetics of male infertility.

### Clinical validity of gene-disease relationships in male infertility

In our literature search, we identified 471 gene-disease relationships that were subjected to critical evaluation. Hereto we used a framework that was designed for interpretation of new research findings in a clinical context in an unbiased way (Smith et al. 2017). The method that we used is a simplified version of the extensive framework used by ClinGen to curate gene-disease relationships and results in similar evidence categories. The method was previously described and proved to be reliable, reproducible and similar to the conclusions of the ClinGen method which makes the method suitable for robust and rapid evaluation of genes in both research and diagnostic sequencing settings (Smith et al. 2017).

The clinical validity of the 471 gene-disease relationships varied widely and ranged from definitive (n=36) to strong (n=12), moderate (n=33), limited (n=86) or no evidence (n=154). A total of 150 gene-disease relationships could not be curated because the tool that we used was not suitable for the type of inheritance pattern observed: we only assessed genes with highly penetrant Mendelian inheritance patterns and excluded mitochondrial and polygenic inheritance patterns (n=45). Furthermore, in several cases the quality of the variant detection was problematic (n=5), essential information like variant information was missing (n=8), or there was insufficient evidence for establishment of the inheritance pattern (n=92) in the original article(s). These results demonstrate that the quality of evidence for gene-disease relationships and reporting of the results varies greatly -a matter that is often not examined or acknowledged in original publications and/or literature reviews.

The curation of all gene-disease relationships was performed with the currently available evidence identified in this literature search. The results are not static and as knowledge increases over time the outcome may be subjected to changes over time. Hence, we expect that a large number of the genes that are currently classified as “Limited”, “No evidence” or “Unable to classify” may still play an important role in male infertility and should therefore not be omitted from future genetic studies.

The number of candidate gene-disease relationships is growing exponentially as a result of the availability of NGS methods and the first half of 2018 has already yielded more novel gene-disease relationships than the full year of 2017 (Figure 2D). However, the number of confidently linked gene-disease relationships is not growing at the same pace. The major reason for this is that most genes have only been found mutated in single patients and functional evidence is lacking. We expect the number of genes confidently linked to azoospermia to grow in the coming years by large-scale data sharing, especially since this is a common form of infertility and genetic components are very likely to play an important role in its etiology (Krausz and Riera-Escamilla 2018).

### Importance of re-evaluation of evidence

The recent availability of large genetic population reference databases facilitates re-evaluation of reported disease-associated variants and allows to determine whether the population frequency of the variant is in line with a reported link to a disorder associated with reduced fitness such as male infertility. Previous reports have shown that healthy participants on average have ~54 exonic variants that were previously reported to be pathogenic, but based on their allele frequency were likely to be misclassified (Lek et al. 2016).

The systematic re-classification of reported genetic sequencing variants in male infertility using this information resulted in some interesting observations. For example, *PICK1* is regularly mentioned as a gene that causes globozoospermia in human patients (De Braekeleer et al. 2015; Ray et al. 2017; Krausz and Riera-Escamilla 2018). However, only one patient with one homozygous variant has ever been described in the initial report of a Chinese globozoospermia patient, and no new patients were published since (Liu, Shi, and Lu 2010). The article was published in 2010, six years before the release of gnomAD which is currently the largest database with allelic information from 123,136 exomes and 15,496 genomes. The variant described in the original publication is present in 1.74% of the East Asian population (http://gnomad.broadinstitute.org/variant/22-38471068-G-A). Calculating the Hardy-Weinberg equilibrium using the incidence of globozoospermia (0.1 % of all male infertility patients) (Holstein et al. 1973) suggests that the maximum allele frequency of this variant can be 0.026%, which is much lower than the observed frequency. Importantly, this maximum allele frequency is still an overestimate as it assumes that all globozoospermia patients are explained by this *PICK1* variant. Although the gnomAD database includes females, may contain male infertility patients and the variant may have reduced penetrance, it is highly unlikely that this particular variant is causing globozoospermia in this patient based on the allele frequency.

Despite the gene-disease relation being based on the wrong data, PICK1 deficiency has been shown to result in disruption of acrosome formation in mice and *PICK1* is expressed in human testis (Xiao et al. 2009). Hence, based on these observations the gene remains an important candidate gene for human male infertility. Similar discrepancies in originally published allele frequencies and currently available allele frequencies were found in several other genes including *NLRP14* (http://gnomad.broadinstitute.org/variant/11-7060977-A-T) (Westerveld et al. 2006), *SEPT12* (http://gnomad.broadinstitute.org/variant/16-4833970-C-T) (Lin et al. 2012) and *RHOXF1* (http://gnomad.broadinstitute.org/variant/X-119243190-C-T) (Borgmann et al. 2016).

Evidence from animal models was often strong and genetic studies clearly benefit from a wealth of studies describing hundreds of well-characterized male infertility mouse models (de Boer, de Vries, and Ramos 2015; Kherraf et al. 2018). However, caution is urged in drawing conclusions about gene function and inheritance mode based on mouse models only. The mouse and human reproductive system are not identical and genes may have (slightly) different functions or transmit disease through different modes of inheritance (Lieschke and Currie 2007). For this reason, we included statistical evidence from large human datasets to supplement the evidence from animal models (Lek et al. 2016; Quinodoz et al. 2017). In case the evidence for inheritance pattern was clearly contradictory between mice and human, we did not evaluate the gene-disease relationship (n=92).

### Recommendations for genetic testing in male infertility

During our study, we noted that international guidelines for nomenclature and interpretation of sequencing variants were often not followed even long after the introduction and world-wide acceptation of these guidelines (den Dunnen et al. 2016; Richards et al. 2015). We identified several errors in nomenclature of sequencing variants and in some cases the variants were not named in a meaningful and unequivocal manner rendering them unusable for assessment. This is in conflict with the FAIR Data Principles for data management and stewardship (Wilkinson et al. 2016). Furthermore, many publications did not mention the expected or proven inheritance pattern or made doubtful conclusions about the mode of inheritance.

In order to ensure efficient sharing and downstream use of newly identified sequencing variants and genes, it is crucial to report variants in an unambiguous and standardized way. In adherence to the standard ACMG guidelines and the best practice guideline of Dutch Genome Diagnostic Laboratories, we have made a list of recommendations for future reporting of novel male infertility variants (Supplemental Table S8). Furthermore, our literature study shows that the quality of evidence of a gene-disease relationship varies greatly. We recommend the use of public and local genomic reference databases, statistical and functional experiments to build evidence for causality (Supplemental Table S9). Due to the sporadic nature of some forms of male infertility, it can be very challenging to acquire multiple patients with variants in the same gene. There are multiple online platforms such as Matchmaker Exchange available for researchers and clinicians which have proven to successfully match patients and uncover rare and novel causes of disease (Philippakis et al. 2015).

### The genetics of human male infertility: overview and future perspectives

Our work shows that the field of genetics of male infertility is rapidly expanding due to the introduction of NGS methods (Figure 2). However, currently of all 471 gene-disease relationships described, only 17% (n=81 gene-disease relationships involving 67 genes) have been at least moderately linked to the disease and an additional 18% (n=86 gene-disease relationships involving 84 genes) are candidate gene-disease relationships with only limited evidence for involvement of the gene in a male infertility phenotype (Table 1, Supplemental Table S4; Figure 3). Caution should be warranted when using genes with limited or no evidence for diagnostic screening.

Similar to other fields in medical genetics, the field of genetics in male infertility has largely focused on inherited variation. Our analysis indicates that 52% of all gene-disease relationships with at least moderate evidence for an involvement in male infertility show an autosomal recessive inheritance pattern (n=42 of 81 gene-disease relationships involving 40 genes). Importantly, many of these genes have been identified in consanguineous families and many of these are associated with very specific and rare sperm defects. It is therefore unlikely that these genes will play a major role in the more common quantitative sperm defects encountered in outbred populations. In contrast, our analysis revealed that only 32% of all gene-disease relationships (n=26 of 81, involving 20 genes) with at least moderate evidence for causing male infertility has an autosomal dominant inheritance pattern, most of which are syndromic presentations.

It may perhaps not be surprising that there is only a limited number of autosomal dominant genes described for male infertility, as pathogenic variation in these genes can only be passed through the maternal line. Importantly, however, studies in intellectual disability and developmental delay have recently pointed to an important role for *de novo* germline mutations resulting in autosomal dominant disease (Vissers, Gilissen, and Veltman 2016). The *de novo* mutation hypothesis for male infertility is further underscored by the fact that *de novo* chromosomal and structural variations are well-known causes of male infertility: Klinefelter syndrome (47,XXY) and AZF deletions almost exclusively occur *de novo* (Lanfranco et al. 2004; Colaco and Modi 2018). The role of *de novo* point mutations, however, remains unexplored in male infertility so far. At the moment, only 3 autosomal dominant genes are moderately linked to isolated male infertility (*DMRT1, KLHL10, SYCP3*). Unfortunately, for none of these genes parental samples were studied to find out whether the variant was paternally or maternally inherited or occurred *de novo*.

### Genetic testing in diagnostic settings

In clinics, genetic testing is offered to infertile men to establish a molecular diagnosis that can be used predict the potential success of fertility treatment options, such as In Vitro Fertilization (IVF), IntraCytoplasmic Sperm Injection (ICSI) combined with TEsticular Sperm Extraction (TESE) or PErcutaneous Sperm Aspiration (PESA) and the risk of transmitting infertility to the next generation. Several studies have shown that men with male factor infertility experience more negative emotional impact such as depressive symptoms, stigma and reduced self-esteem, than men whose partners were infertile or men of couples diagnosed with unexplained infertility (Fisher and Hammarberg 2012). The standard use of NGS in the diagnostic work-up of male infertility could lead to more men receiving a diagnosis, or explanation, and therefore possibly influence this emotional burden in a positive way.

The recommendations for genetic testing during the diagnostic work-up of male infertility have only minimally changed over the last 20 years and most of these recommendations still focus on the well-known and common causes of male infertility that were already known in the 1990’s (Barratt et al. 2017; Jungwirth 2018). For cost-efficiency, there are guidelines to help stratify patient groups to receive pre-conceptive genetic tests such as karyotype analysis, AZF deletion tests or a screening for pathogenic variants in a single gene involved in a specific phenotype such as CBAVD or Kallmann syndrome. A recent World Health Organization study on the diagnosis on male infertility suggested to at least perform karyotyping and AZF deletion tests in men with non-obstructive azoospermia or extreme oligoasthenoteratozoospermia (OAT) without a history of a known cause of spermatogenic failure such as chemotherapy, varicocele, orchitis or bilateral cryptorchidism (Barratt et al. 2017). However, after stratification, in approximately 40% of all male infertility patients no genetic cause is found with the above mentioned tests (Krausz and Riera-Escamilla 2018) and this strongly suggests that much more genetic research is required and at the same time the use of other diagnostic assays should be considered.

Testing all patients for all genetic anomalies was very costly for a long time. However, in light of the recent developments of novel sequencing technologies, it is now possible to consolidate one or multiple tests in a single NGS assay which will help to cut the costs. The first examples of NGS-based screening methods have been described for male infertility (Oud et al. 2017; Fakhro et al. 2018; Patel et al. 2018). The European Society of Human Genetics (ESHG) and the European Society for Human Reproduction and Embryology (ESHRE) have recently made a recommendation for developing and introducing new tests, specifically for extended carrier screening (Harper et al. 2018). Genetic tests should be designed to achieve high clinical validity, establish clinical utility, minimize secondary findings such as carriership of cancer-predisposing variants as the capacity of follow-up counseling is limited and furthermore they emphasized that providers should take into account that there are inter- and intra-population and individual differences for genetic risk and disease. For example, when considering the effect of immigration of non-European populations on counseling and interpretation of uncommon disease-associated variants (Harper et al. 2018).

With the current rise of NGS in the field of male infertility, the number of novel pathogenic variants and genes will grow rapidly (Figure 2). The identification of novel disease genes allows for selection of genes for male infertility gene panels. For diagnostic purposes, gene panels should contain genes with a minimal level of evidence of involvement with disease. We recommend to include genes with an evidence classification of at least “Moderate” for the composition of diagnostic gene panels. We recommend the inclusion of genes involved in syndromic forms of male infertility. The severity of several syndromes relies on the damaging effect of the mutation(s) and the spectrum may span combinations of recognizable associated features to isolated infertility. It is therefore possible that patients referred to for infertility suffer from a mild syndrome which features were missed or not present upon anamnesis.

### Conclusion

In this clinical validity assessment, we evaluated a total of 471 gene-disease relationships involving 438 genes with reported monogenic association to male infertility and identified 81 gene-disease relationships with at least moderate evidence for a role in male infertility. Both our results as well as our objective approach and recommendations may aid the robust and rapid identification and incorporation of novel genes in male infertility diagnostics.

## Author roles

M.S.O., L.V., L.E.L.M.V, and J.A.V. designed this study and L.R., L.E.L.M.V. and J.A.V. supervised this study. M.S.O. and L.V. selected studies for the inclusion and evaluation of quality. M.S.O. evaluated the quality of all included publications and L.V. reviewed and verified the results. Disagreements in the inclusion and evaluation process were solved by consensus between all authors. All authors made substantial contributions to the interpretations of the results. M.S.O. and L.V. prepared the figures and M.S.O., L.V. and R.M.S. wrote the first draft of the manuscript. All authors contributed to the revision process.

## Acknowledgements

This work was supported by The Netherlands Organisation for Scientific Research. The authors would like to thank the members of the International Male Infertility Genomics Consortium for their constructive comments.

